# Human pluripotent stem cell-derived bronchial airway organoids provide insights into differential innate immune and long-term responses to SARS-CoV-2 infection in healthy and COPD

**DOI:** 10.1101/2025.06.30.662440

**Authors:** Lisa Morichon, Jitendriya Swain, Nathalie Gros, Amel Nasri, Florent Foisset, Gaetan Galisot, Victor Racine, Saïd Assou, Arnaud Bourdin, John De Vos, Delphine Muriaux

**Author notes:** **Correspondence:** Co-last senior authors, (DM); (JDV).

## Abstract

Respiratory infections are a major global health concern, as underscored by the COVID-19 pandemic. To better understand bronchial tissue responses to viral infection, we have developed a preclinical in vitro model mimicking the multiciliated airway epithelium, from induced pluripotent stem cell (iPSC) and cultured in an air-liquid interface (iALI). By using iPSCs reprogrammed from patients with chronic obstructive pulmonary disease (COPD), we successfully generated a fully differentiated and functional bronchial epithelium exhibiting key COPD features with goblet and basal cell hyperplasia and tissue inflammation. SARS-CoV-2 could infected and replicated for several weeks in both healthy and COPD models, with a recurrent peak at 3 days after infection. Infected iALI exhibited cilia destruction and increased mucus secretion. Innate immune response of different infected iALI reveals a differential expression of interferon-stimulated genes (ISGs) and pro-inflammatory cytokine secretion. Notably, COPD iALI displayed an earlier innate immune response to SARS-CoV-2 infection as compared to healthy iALI, suggesting a genetic susceptibility of COPD iALI towards inflammation induced by SARS-CoV-2 infection, and a less efficient response to antivirals. In conclusion, our study demonstrates that the iALI bronchial organoid model is a powerful tool for investigating bronchial tissue responses to long term respiratory viral infections, antivirals, and patients with COPD or other airway pathology.

**Grapical Abstract:** 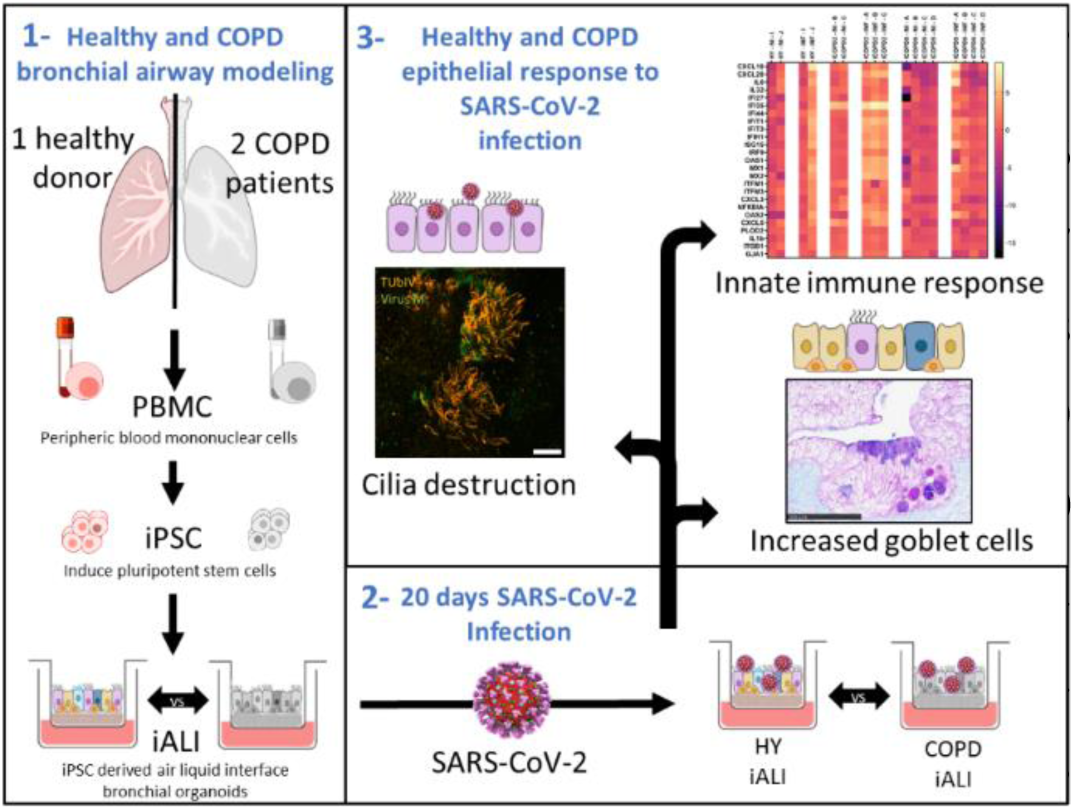

**HighLights:**

- hiPSC-derived COPD airway organoids
- SARS-CoV-2 productively infects induced pluripotent stem-cell derived bronchial organoids iALI and persisted over the long term
- SARS-CoV-2 iALI infection results in cilia destruction, increased mucus secretion and a strong innate immune response
- SARS-CoV-2 infection elicited a higher and earlier innate immune response in iCOPD
- SARS-CoV-2 infection in iCOPD respond less to antivirals

**Short Abstract:** SARS-CoV-2 causes severe lower respiratory tract infection in COVID-19 patients, which can persist over time. Here, we used an in-house developed in vitro airway organoid derived from induced human pluripotent stem cells (iALI) to study SARS-CoV-2 infection over long term in healthy or COPD patients whom respiratory failure is at risk during infection. Our results show that SARS-CoV-2 infection results in high and lethal infection of bronchial epithelial cells, that persist over time, inducing mucus secretion, destruction of ciliated cells and specific cytokine release. A late innate immune response is observed in the healthy iALI, while in iCOPD, it appears earlier and stronger, suggesting a different sensing of SARS-CoV-2 in COPD patients, accompanied by a reduce sensitivity to antivirals. In conclusion, our study demonstrates that the iALI organoid model is a powerful tool for investigating bronchial tissue responses to long term respiratory viral infections, from healthy to pathologic patients.

## Introduction

Chronic obstructive pulmonary disease (COPD) is the fourth leading cause of death worldwide, accounting for 3.5 million deaths in 2021, approximately 5% of all global deaths (WHO, November 4^th^ 2024). Primarily caused by exposure to harmful particles, particularly cigarette smoke in westernized countries, COPD is characterized by chronic bronchitis and emphysema (1–3). This disease starts and progresses at the small airway level narrowed by thickened walls, excessive mucus secretion and chronic inflammation, worsened by loss of alveolar attachments Recent studies have shown that COPD patients have a fourfold increased risk for severe clinical outcomes — including hospitalization, ICU admission, and mortality— when infected with COVID-19 (4–7). This heightened susceptibility may be attributed to increased ACE2 expression in the lungs of smokers and COPD patients (8–10), impaired muco-ciliary clearance (MCC) (2,11–14) and/or dysregulated immunity (1,2,15,16), although controversies persist. Traditionally, COPD research has relied on blood samples (17), lobectomies (18), or murine models (19,20). However, the recent development of lung organoids models has significantly expanded research possibilities. Since COVID-19 pandemic, airway epithelium models have also been used intensively to investigate SARS-CoV-2 pathophysiology (12,21–23)

In vitro airway models are created using progenitors derived either from primary samples of the upper (nasal or tracheal) or lower (bronchial or alveolar) airways, or by differentiation of induced pluripotent stem cells (iPSCs). The advantage of using primary progenitor cells lies in their origin from adult lungs, which retain both the genetic and epigenetic traces of each individual (24,25). This allows mature epithelia to be obtained and pathological phenotypes to be reproduced (10,26), but also lead to high interindividual variability in results. Moreover, primary models are expensive and limited in quantity. In contrast, the process of reprogramming somatic cells into iPSCs, from blood mononuclear cells (PBMCs) for example (27), eliminates epigenetic marks (28,29), and allows them to be amplified in cell culture almost without limit. It is therefore possible to genetically modify iPSCs and reproduce all the experiments required for a project from a single donor. In addition, iPSCs differentiation produces both airway epithelium and mesenchyme, which provides physiological support for the epithelium and improves its survival (30) through time and aggressions. However, the epithelium obtained is less mature than those derived from primary cells (31–33).

Few models derived from primary progenitors were used to investigate the response of COPD to SARS-CoV-2 infection (10,34–36). The first model involved an air liquid interface (ALI) culture of COPD human bronchial epithelial cells (NHBE), which successfully reproduced COPD phenotypes, including basal and goblet cell hyperplasia and squamous metaplasia. This model also demonstrated a high susceptibility of goblet cells to SARS-CoV-2 infection (34). A second study used NHBE and human nasopharyngeal epithelial cells and differentiated them in apical out airway organoids (10). These models also exhibited characteristics of the COPD phenotype, including goblet cells hyperplasia and reduced ciliary beat frequency. Both studies showed higher SARS-CoV-2 replication rates in COPD airway organoids, exacerbating the pathophysiology of COPD. In contrary, recent study demonstrated lower SARS-CoV-2 infection and inflammatory response in COPD human airway epithelia (HAE) as compared to healthy HAE, associated with protective characteristics of COPD HAE (35). It is important to note, that all these studies focus on early response to infection and did not investigate tissue response on the long-term after infection. Overall, the available results still lack comparative data and investigation of long-term infection tissue impact.

Here we propose another in vitro preclinical model, called iALI, derived from human induced pluripotent stem cells (iPSC) and cultured at the air-liquid interface (ALI), named iALI, to study COPD and SARS-CoV-2 infection (30). Working on iPSC-derived bronchial epithelium obtained from healthy and COPD primary blood mononucleated cells, we demonstrated successful persistent infection of both healthy and COPD iALIs for up to 21 days. We highlight tissue response to infection, as well as the epithelium innate immune response. Our results show that the iALI model is relevant to study the COPD pathology, SARS-CoV-2 infection over the long term, antiviral drug testing as pre-clinical models and it opens the possibility to target molecular pathways and explore the long-term impact of COVID-19 in the airways.

## RESULTS

### Characterization of the bronchial epithelium obtained from differentiation of Healthy and COPD iPSC-derived iALI organoids

Multi-ciliated bronchial epithelium were established by methods previously reported (30) from 3 different iPSC cell lines derived from 1 healthy donor (HY03) and 2 very severe and early COPD patients (iCOPD2, iCOPD9) (Fig. 1A) (27,37). HES staining of paraffin embedded cross section confirmed proper morphology of iALI with pseudostratified cylindrical bronchial epithelium, cilia at the apical side and under the basal membrane a thick layer of mesenchyme sometimes including cartilage (Fig. 1B, Supplemental Fig.1). Immunofluorescence staining of cross-section demonstrated presence of ciliated cells (*CDHR3*), goblet cells (*MUC5AC*), basal cells (*KTR5, P63*) and club cells (*SCGB1A1*) (Fig. 1C-E). Quantification of gene expression in bronchial epithelial cell types was done by RTqPCR. Ciliated cells levels were evaluated by *FOXJ1* and *CCDC40* relative gene expression (Fig. 2A), goblet cells with *MUC5AC* and *MUC5B* (Fig. 2B), basal cells with *KRT5* and *P63* (Fig. 2C), neuroendocrine cells with *CHGA* (Fig. 2D) and club cells with *SCGB1A1* (Fig. 2E). Our results validate proper differentiation of iPSc into main different cell types of the bronchial airway epithelium.

**Figure 1:**
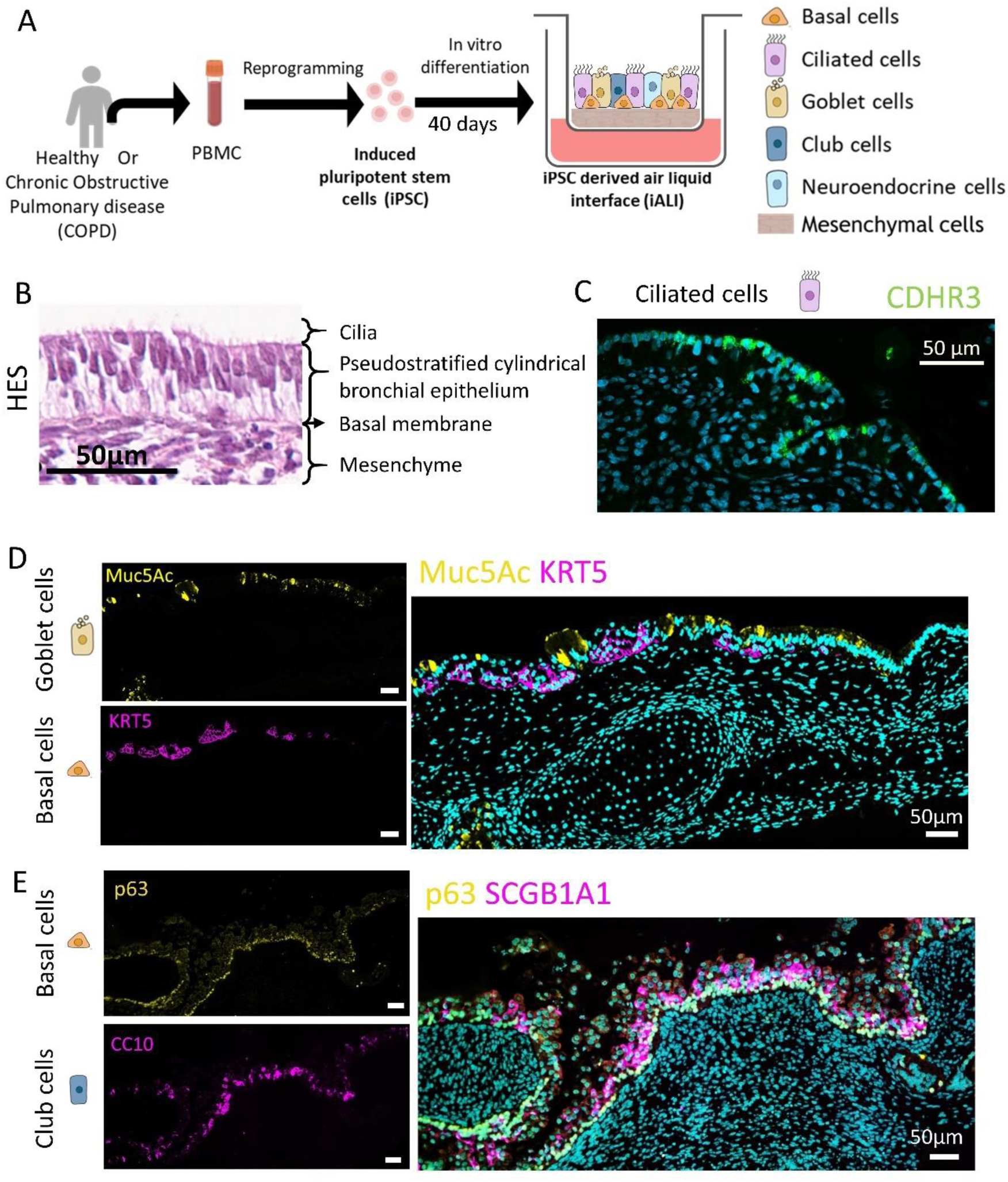
Bronchial epithelium imaging characterization of iALI organoids. (A) Schematic representation of iALI production protocol and epithelial cells composition. (B) iALI epithelial morphology was confirmed by hematoxylin eosin safran (HES) staining. (C-E) The epithelial cells presence was evaluated by immunofluorescence (IF) of iALI cross sections. Representative IF images of iALI stained for ciliated cells (green, CDHR3 : C), goblet cells (yellow, Muc5Ac : D), basal cells (pink, KRT5 : D; yellow, p63 : E) and club cells (pink, SCGB1A1: E). Scale bar is 50µm.

**Figure 2:**
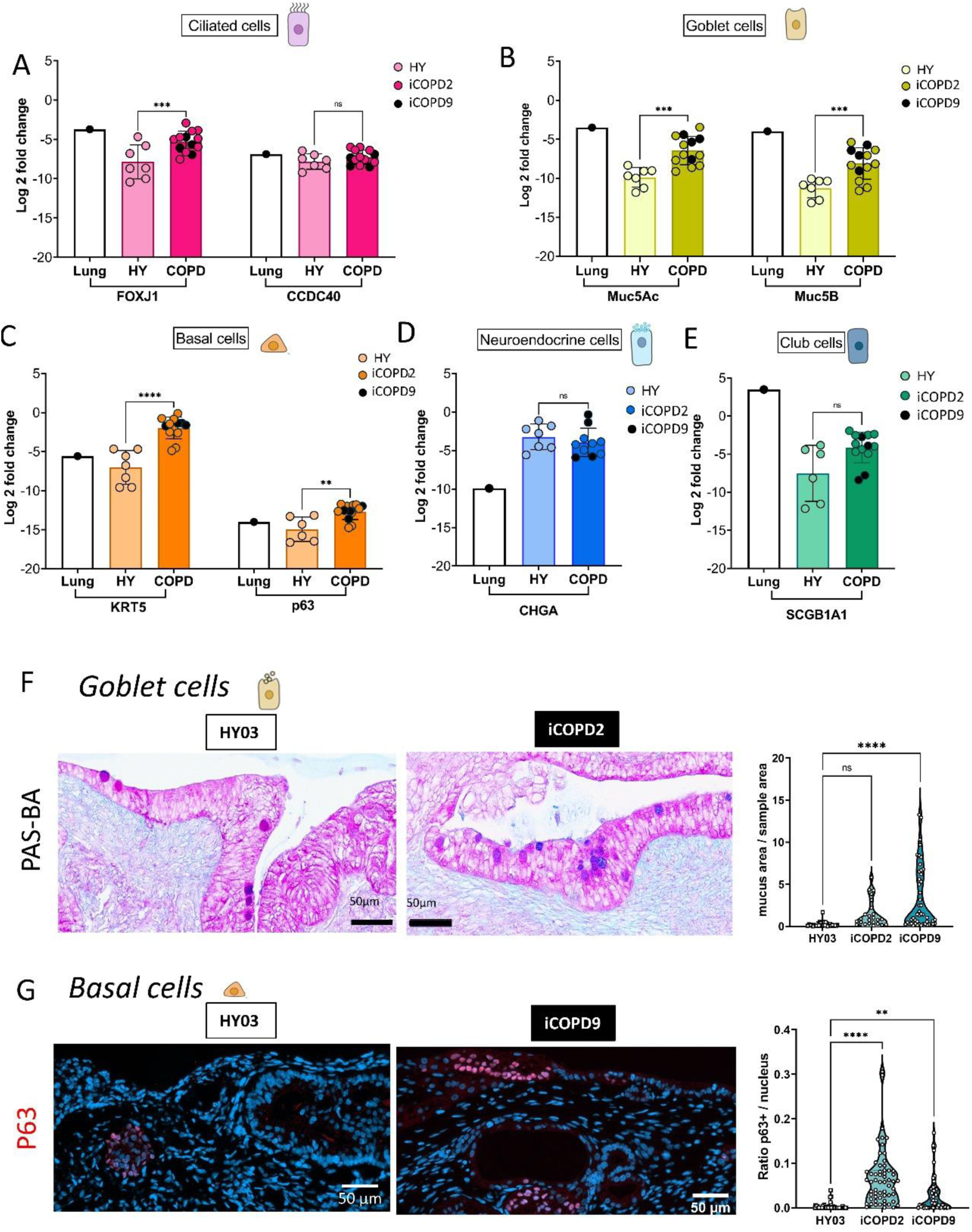
Comparison of the cellular composition of iALI organoids derived from healthy and COPD patients. (A-E) Epithelial cells presence in non-infected HY03 (N=5, n=7), iCOPD2 (N=2, n= 4) and iCOPD9 (N=4, n=9) iALI was confirmed with the measure of gene expression relative to GAPDH by RTqPCR. Human lung total RNA was used as a positive control to measure the expression of ciliated cells (*FOXJ1, CCDC40*: A), goblet cells (*MUC5AC, MUC5B* : B), basal cells (*KRT5, P63* : C), club cells (*SCGB1A1* : D) and neuroendocrine cells (*CHGA* : E). (F) Periodic acidic Schiff and blue alcian (PAS-BA) staining of HY03 (N=2, n=6), iCOPD2 (N=1, n=4) and iCODP9 (N=2, n=8) iALI allowed quantification of goblet cells (dark pink and purple). (G) Quantitative analysis of basal cells levels (red, P63) in HY03 (n=2), iCOPD2 (n=2) and iCOPD9 (n=2) iALI. (A-G) *p < 0.05, **p < 0.01, ***p < 0.001, **** p < 0.0001, NS represents not significant, (A-E) two-way ANOVA with Sidak multiple comparisons test. (F-G) one-way ANOVA with Dunnett’s multiple comparisons post-test. N= Number of experiments, n= number of samples analyzed.

### The bronchial airway epithelium differentiated from COPD iPSC mimics the characteristic features of COPD

Chronic obstructive pulmonary diseases (COPD) is known for impaired epithelial phenotype including goblet cells hyperplasia, basal cell hyperplasia and inflammation (2,16,34,38–40). RTqPCR analysis of iALI revealed a significant increased relative expression of goblet cell markers (*MUC5AC* and *MUC5B*) and basal cell markers (*KRT5, P63*) in the COPD iALI compared to healthy iALI (Fig. 2 B, C). Goblet cell hypertrophy was confirmed by analysis of PAS-BA stained cross section that shows a significant increase in iCOPD9 iALI and a tendency in iCOPD2 iALI (Fig. 2F). Quantification of P63 positively stained nucleus validated a significant increase of basal cells levels both in iCOPD2 and iCOPD9 iALI (Fig. 2G). Inflammation was evaluated by measuring relative expression of innate immune response pathway genes. At steady state, *CXCL5*, *CXCL10*, *IFI35* and *OAS2* were significantly increased in iCOPD2 samples. No difference was found in iCOPD9 samples at a non-stimulated state (Supplemental Fig.2). Cytokine expression levels were also measured by flow immunoassay and revealed no difference at rest between healthy and COPD iALI (Supplemental Fig.3). So, iALI derived from COPD iPSc reproduces COPD characteristics, that are signed by increased mucus, basal cells and latent inflammatory state. These results suggest either a COPD’s genetic susceptibility present in our 2 COPD donors or maintenance of some epigenetic features despite reprogramming process of iPSC.

### Bronchial airway epithelial cells of iALI, particularly ciliated cells, are infected by SARS-CoV-2

The air-liquid interface culture associates with the polarization of the tissue and set epithelial cells at the apical side of the iALI and in some invaginations deeper in the tissue (Supplemental Fig. 1 & 4B). So, susceptibility of iALI to infection by the SARS-CoV-2 Delta variant was tested by adding viral dilutions at the apical side (Fig. 3A) at an estimated multiplicity of infection (MOI) of 0.05 (2,5.10^5 PFU/well). Proper infection of iALI was confirmed by positive staining of SARS-CoV-2 membrane in healthy iALI cross section 3 and 11 dpi (Fig. 3B). Virus was detected from day 1 to day 11 in airway epithelial cells and later in mucus plugs (Supplemental Fig. 4A). Quantification of the infection levels was done by RTqPCR analysis of SARS-CoV-2 gene E relative expression in iALI cell lysates at several time points post infection (Fig. 3C).). While we noticed some variability between experiments, a significant decrease of the viral replication in the iCOPD9 iALI compared to HY03 iALI was observed at 1 dpi, and at 3, 4 and 6 dpi compared to iCOPD2 iALI. Despite a decrease at 1 dpi, viral replication was significantly increased 6 dpi in iCOPD2 iALI compared to HY03. Viral release was further investigated in everyday apical washes by RTqPCR measurements of gene E absolute concentration (Fig. 3D) and by plaque assay for the infectious titer (Fig. 3E & F). In all samples, viral production peaked at 2 dpi and then decreased but maintained itself for up to 21 dpi. Except for a lower gene E absolute concentration in iCOPD9 iALI at 2 dpi, compared to HY03, no significant difference was found between HY03, iCOPD2 and iCOPD9 iALI. Moreover, ACE2 and TMPRSS2 relative gene expressions were similar in HY03, iCOPD2 and iCOPD9 non-infected and infected iALI (Supplemental Fig.1B & C) indicating that virus entry/infection should be similar between samples. Lastly, SARS-CoV-2 infection and replication in healthy and iCOPD2 iALI was inhibited by the antiviral Remdesivir drug targeting the SARS-CoV-2 RNA dependent RNA polymerase, confirming the infection of the iALI by the virus (Fig.3 G, H, I). The inhibitor of the viral replicase reduced infectivity in both healthy and COPD iALI. Surprisingly, we notice a lower efficiency of the Remdesivir in iCOPD2 than in healthy iALI with a viral log10 reduction of respectively 1,58 and 2,16 at 2 days post infection respectively (Fig. 3 G-I). This difference could be caused by a different diffusion of the antiviral drug in COPD iALI with an impact of the increased mucus in pathological iCOPD organoids. Overall, our data suggest a susceptibility of HY03 and COPD iALI to SARS-CoV-2 infection albeit differences between donors.

**Figure 3:**
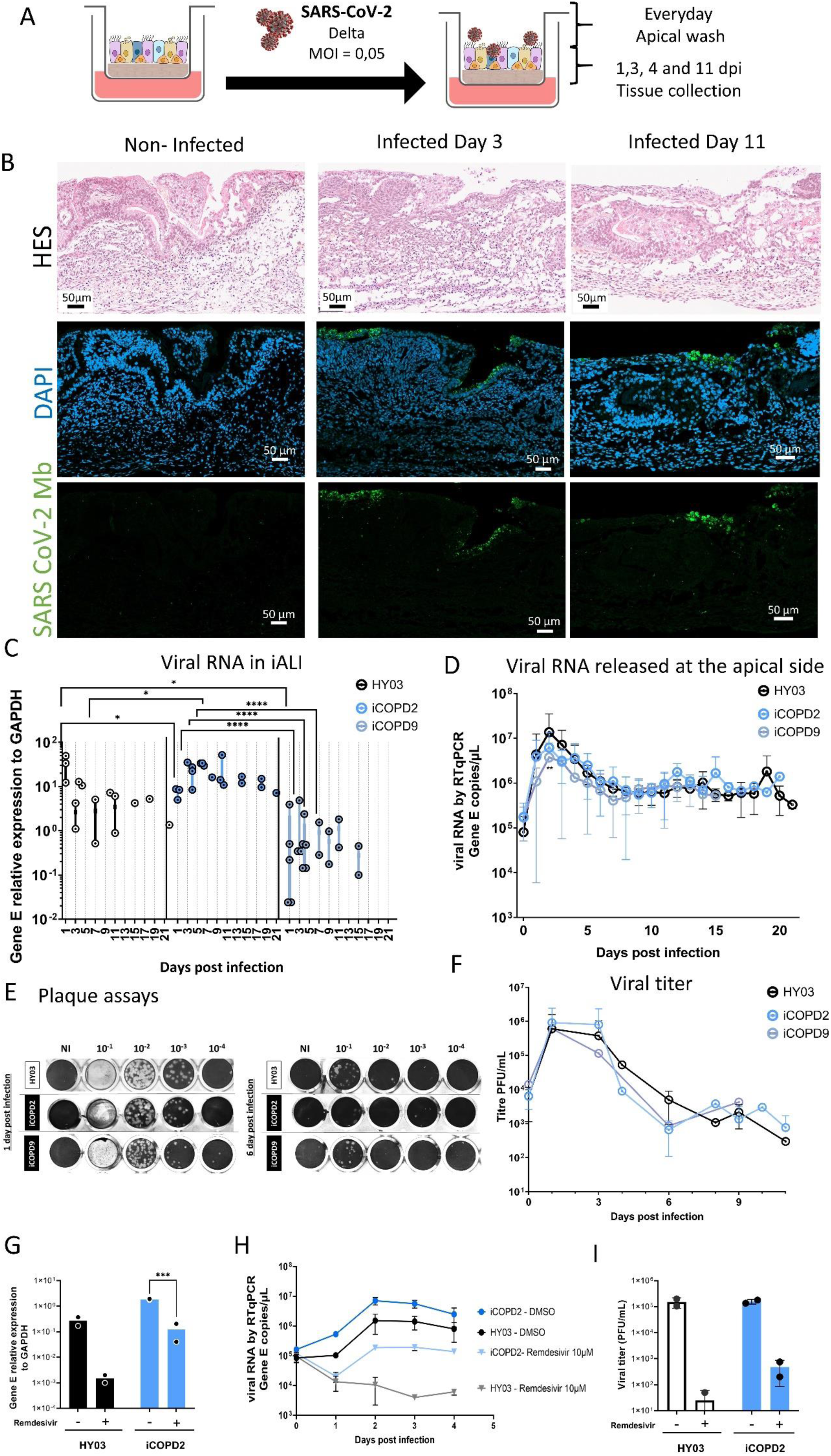
SARS-CoV-2 infection in Healthy and COPD iALI organoids. (A) Schematic representation of iALI infection and collection parameters. (B) SARS-CoV-2 infection was visualized by HES staining and labelling for SARS-CoV-2 membrane (Virus M, green) healthy non-infected and infected (3dpi &11dpi) iALI cross section. (C) RT-qPCR analysis of viral envelope gene relative expression compared to GAPDH was done in the HY03 (N=4), iCOPD2 (N=3) and iCOPD9 (N=5) infected iALI cells. (D) RT-qPCR quantifications of apical wash viral envelope gene on HY03 (N=5, n=9), iCOPD2 (N=4, n=4), iCOPD9 (N=4, n=7). (E-F) Viral titer was measured by plaque assay in apical wash of HY03 (N=3), iCOPD2 (N=4), iCOPD9 (N=4). Representative images of plaque assays at 1- and 6-days post infection (E). (G-I) HY03 and iCOPD2 iALI were treated with DMSO (0,1%) or Remdesivir (10 µM) 2 hours prior infection by SARS-CoV-2 (Delta, MOI = 0,05). (G) Viral RNA (gene E) relative expression to GAPDH was measured by RT-qPCR in iALI cell lysates 96hpi (H) Quantification by RTqPCR of viral RNA (gene E) released every day post infection at the apical side of treated iALI is presented as gene E concentration (I) Viral particles released at the apical 2dpi were quantify by plaque assays.(C-I) *p < 0.05, **p < 0.01, ***p < 0.001, **** p < 0.0001, two-way ANOVA with Sidak’s (C, G) or Tukey’s (D & F) multiple comparisons post-test. N= Number of experiments, n= number of samples analyzed.

### SARS-CoV-2 infection impact on healthy and COPD bronchial airway organoids

The impact of SARS-CoV-2 on the iALI tissue was first visualized and quantified on HES labelled cross section images of infected samples, where we noticed a decrease in the relative epithelium area, especially the apical one (Supplemental Fig. 4 B, C). Further consequence on the epithelium was investigated by RTqPCR measurement of cell type marker relative expression in infected iALI compared to non-infected (Fig. 4). While no significant difference was found, we see a tendency of increase in goblet cell and basal cell markers in both HY03 and COPD infected iALI. Mucus cells were also quantified by PAS-BA labelled cross sections(Fig. 5 A & B). Results show a significant increased amount of goblet cells in healthy iALI 3 dpi and 11 dpi. As already described, goblet cell quantity and area in iCOPD is significantly higher than in HY03 iALI (Fig. 2B &F). Yet, it still tends to increase in infected samples at 1 and 11 dpi (Fig. 5B). It is interesting to note that mucus level in healthy iALI at 11dpi and in iCOPD at 1dpi are similar, suggesting that iCOPD have a strongly stimulated basal level. We then focused on the impact of SARS-CoV-2 infection on ciliated cells (Fig.5 B-D). We labelled cilia (TUBIV) and SARS-CoV-2 membrane (Virus M) in healthy and COPD, non-infected and infected (3dpi) iALIs (Fig. 5C). High resolution fluorescence microscopy imaging confirmed infection, particularly in ciliated cells, showing a destruction of the cilia by the virus in ciliated cells where the virus was abundant. Ciliated cells quantification show a significant decrease of ciliated cells in infected samples, by 25% in HY03 iALI and 30% in iCOPD iALI (Fig. 5 C, D). Thus, infected iALI organoids reproduces the consequence of infection observed on bronchial epithelium derived from primary cells (10, 34, 35), characterized by an increase of mucus secretion and a destruction of cilia cells (Fig. 5). Despite differences between infected and non-infected samples, we see no apparent difference between HY03 and COPD epithelial responses to SARS-CoV-2 infection.

**Figure 4.**
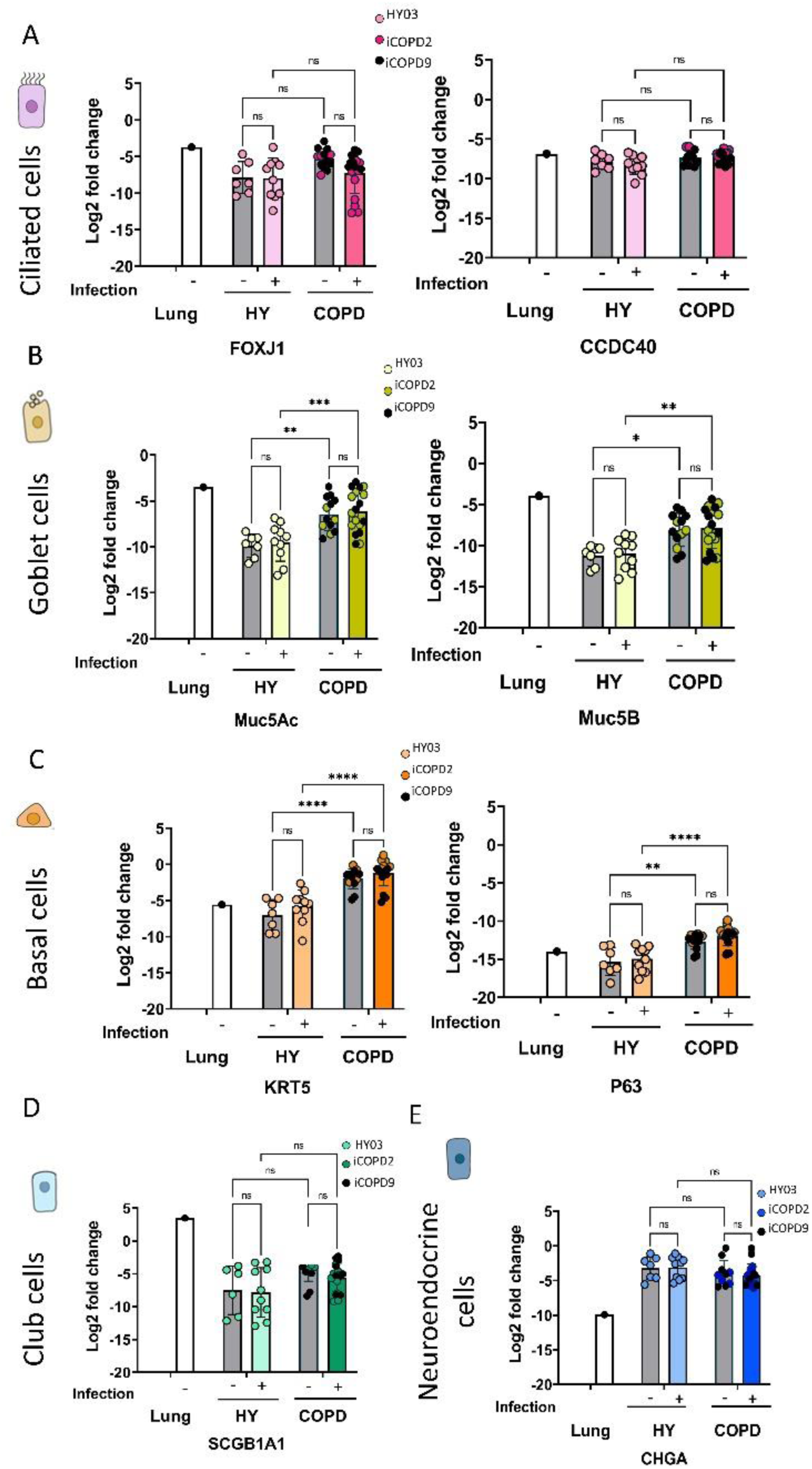
: Impact of SARS-CoV-2 infection on the cellular composition of healthy and COPD iALI organoids. (A-E) Epithelial cells presence in non-infected HY03 (N=5, n=7), iCOPD2 (N=2, n= 4) and iCOPD9 (N=4, n=9) iALI and SARS-CoV2 infected HY03 (N=6, n=10), iCOPD2 (N=3, n= 9) and iCOPD9 (N=4, n=9) iALI was confirmed with the measure of gene expression relative to GAPDH by RTqPCR. Human lung total RNA was used as a positive control to measure the expression of ciliated cells (*FOXJ1, CCDC40*: A), goblet cells (*MUC5AC, MUC5B* : B), basal cells (*KRT5, P63* : C), club cells (*SCGB1A1* : D) and neuroendocrine cells (*CHGA* : E). *p < 0.05, **p < 0.01, ***p < 0.001, **** p < 0.0001, NS represents not significant, (A-E) two-way ANOVA with Tukey’s multiple comparisons test.

**Figure 5:**
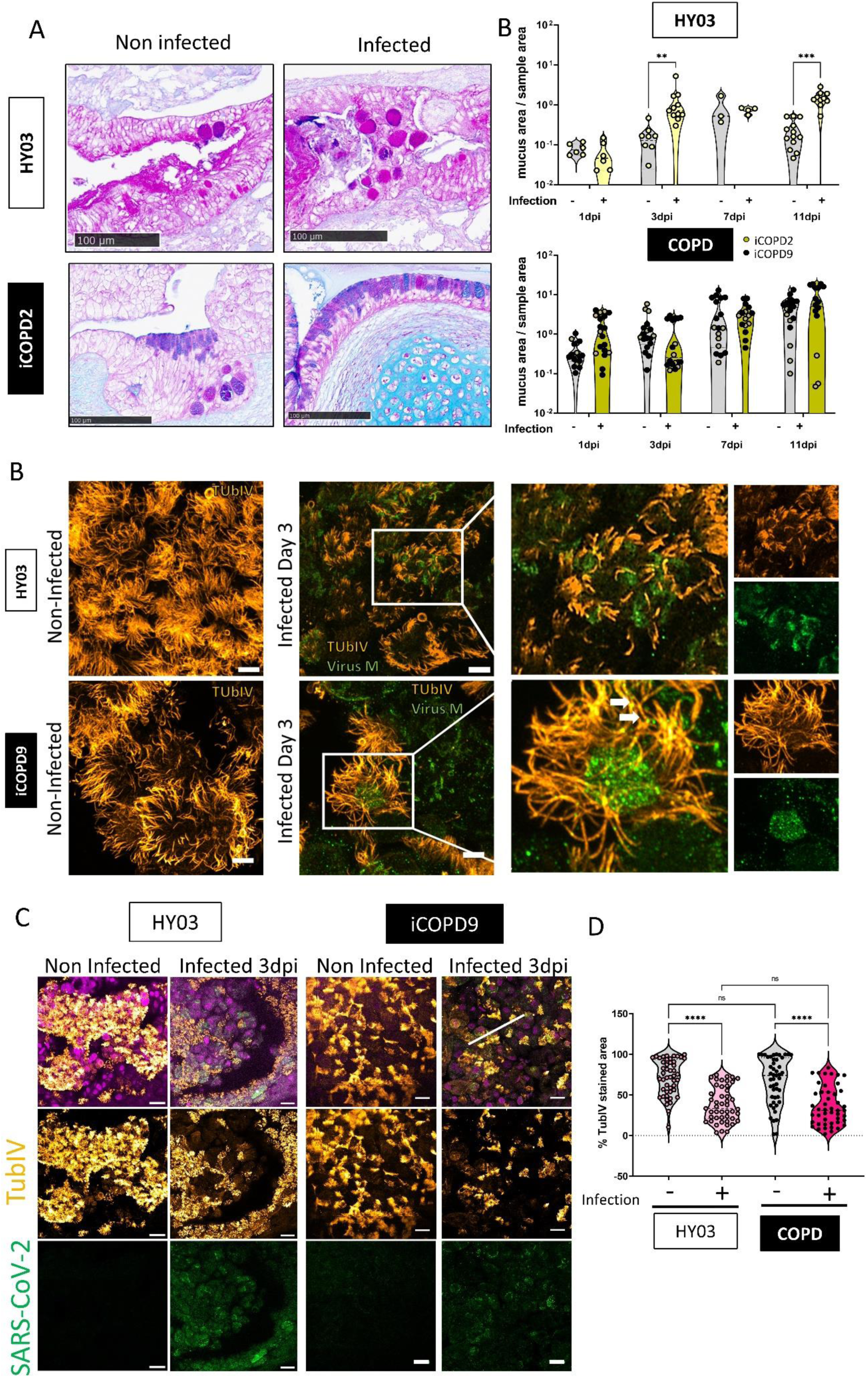
Impact of SARS-CoV-2 infection on HY03 and COPD iALI organoids. (A) Representative imaging of periodic acidic schiff and Blue acian (PAS-BA) staining of non-infected and infected HY03 and iCOPD2 iALI. (B) Quantification of goblet cells (dark pink and purple) normalized to full sample area on non-infected and infected HY03 iALI (N=2, n=6), iCOPD2 (N=1, n=4) and iCODP9 (N=2, n=8). (B) High resolutions images confirmed infections by staining SARS-CoV-2 (green, Virus membrane) and cilia (orange, TubIV) on non-infected and infected (3dpi) healthy and iCOPD2 iALI. Scale bar = 5µm. (C) High resolution 3D images of non-infected and infected (3dpi) HY03 and iCOPD9 iALI stained for ciliated cell (Yellow, Tub IV), Virus (Virus membrane, green) (D) Quantifications of the percentage of TubIV-positively stained area for both infected and non-infected (3dpi) HY03 and iCOPD2 on 47 regions of interest (25µm x25µm). Representative images, scale bar = 100µm (A) 5µM (B) and 50µm (C). *p < 0.05, **p < 0.01, ***p < 0.001, **** p < 0.0001, NS represents not significant, one-way ANOVA with Tukey’s multiple comparisons post-test (B), two-way ANOVA with sidak multiple comparisons post-test (D). N= Number of experiments, n= total number of samples analyzed.

### iCOPD organoids have an impaired innate immune response to SARS-CoV-2 infection

Innate immune response to infection was evaluated by measuring the expression of 25 common interferon-stimulated (ISG) using RTqPCR in HY03, iCOPD2 and iCOPD9 iALI at 1, 4, 11 and 18 dpi (Fig. 6 & Supplemental Fig. 2 & 5). At early time points in HY03 iALI, increased expression of only few of these genes was observed in infected conditions with only *IFIT1*, *ISG15* and *OAS2* significantly higher at 1 and 4 dpi, plus *MX1* at 4 dpi. *IFIT1* and *ISG15* gene expressions were also always found significantly increased at 1, 4 and 11 dpi for both COPD iALI, as well as several others. For iCOPD2, we also found higher expressions of *IFIH1*, *MX1*, *MX2*, *ITFM3* and *OAS2* at 1 dpi and of *MX1* and *CXCL10* for iCOPD9. At 4 dpi for both COPD iALI, there was an overexpression of *IFI35* and *ITFM3*. Infected iCOPD9 iALI presents a significant increased expression of these following genes at 4 dpi: *CXCL10*, *IFI44*, *IFIT3*, *IFIH1*, *ISG15*, *MX1*, *MX2*, *OAS2* and *ITFM1*. At 11 and 18dpi, we compared innate immune response in HY03 and iCOPD2 iALI. At 11 dpi, *IFIT1*, *ISG15* and *OAS1* were commonly significantly overexpressed. While iCOPD2 had a persistent significantly increased expression of only *IFI35*, surprisingly HY03 iALI showed a significant over expression of several interferon pathway genes: *IFI27, IFIT3, MX1, MX2* and *OAS2*. At 18 dpi, although not statistically significant, we keep a similar pattern than at 11 dpi (Supplemental Fig. 5).

**Figure 6:**
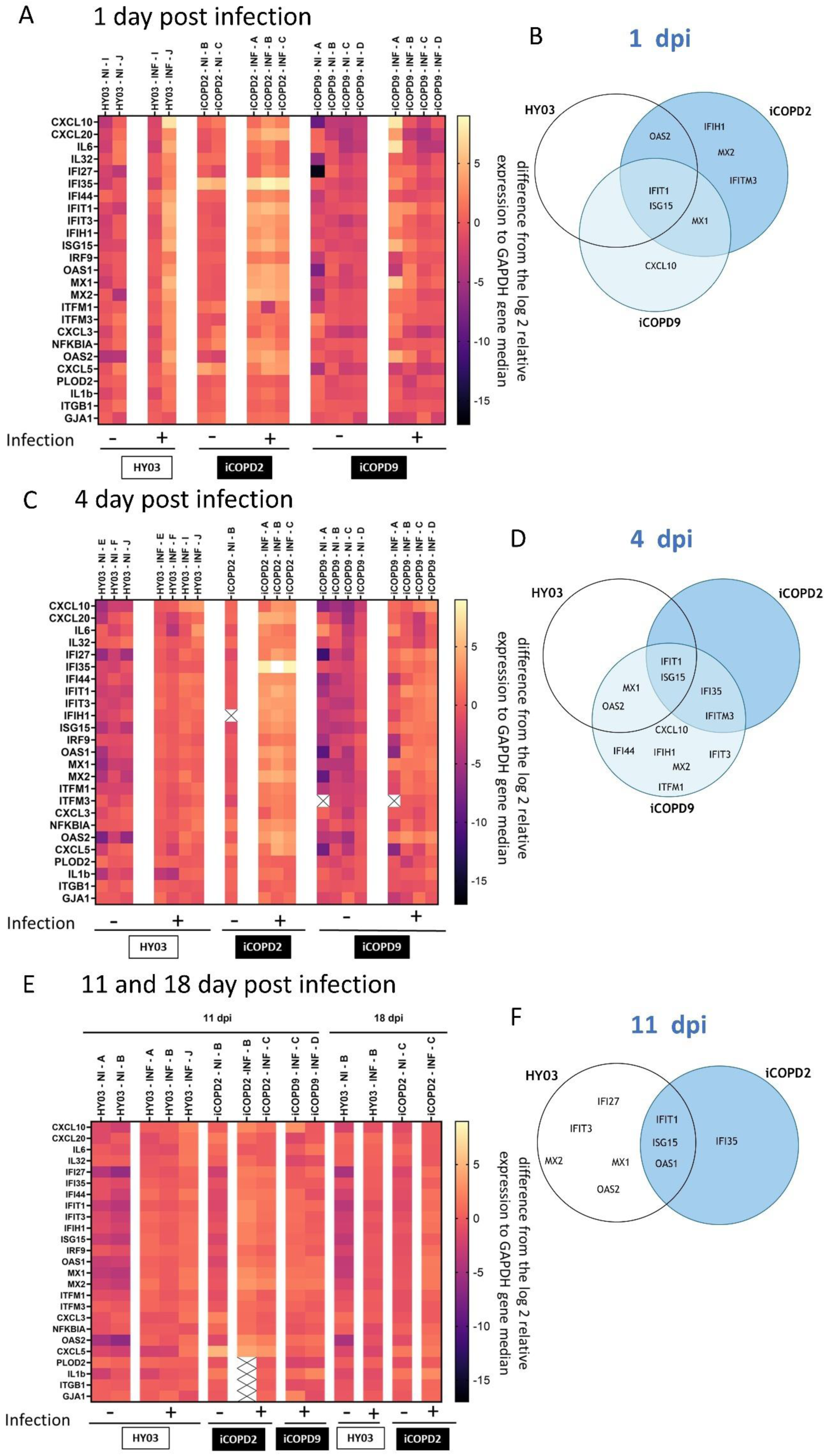
Transcriptomic analysis of innate immune response on infected HY03 and COPD iALI. (A) 1dpi (C) 4dpi (E) 11dpi innate immune response gene expression levels measured on non-infected (NI) and infected (INF) HY03, iCOPD2 and iCOPD9 iALI, by the relative expression of *CXCL10, CXCL20, IL6, IL32, IFI27, IFI35, IFI44, IFIT1, IFIT3, IFIH1, ISG15, IRF9, OAS1, MX1, MX2, ITFM1, ITFM3, CXCL3, NFKBIA, OAS2, CXCL5, PLOD2, IL1b*. Integrity of the tissue is shown by ITGB1 and GJA1 relative expression. Each value represents the difference from the log2 relative expression to GAPDH gene median and is depicted according to the color scale shown at the right (−15 to 5). Each experiment is coded with an alphabet letter (ie., A to J). (B) 1 dpi, (D) 4 dpi and (F) 11dpi significantly overexpressed gene in infected compared to non-infected samples in one or several organoids cell line are represented in a Venn diagram.

Our results show a reproducible heterogeneous pattern of the response to the SARS-CoV-2 infection between iCOPD9 and iCOPD2 iALI organoids overtime, suggesting that COPD patients do not respond in an equivalent manner. It is also important to note that expression levels detected in infected iCOPD2 samples were often significantly higher than for HY03 and iCOPD9 iALI, for instance: *CXCL20*, *IFI35*, *IFIH1*, *ISG15*, *CXCL5*, *ITFM3* and *NFKBIA* (Supplemental Fig. 2). In summary, innate immune response to SARS-CoV-2 infection was more important at 1 dpi for iCOPD2, at 4dpi for iCOPD9 and 11dpi for HY03 iALI. *IFIT1* and *ISG15* were overexpressed in every iALI at 1, 4 and 11 dpi. *MX1*, *MX2* and *OAS2* were successively overexpressed in iCOPD2 at 1 dpi, iCOPD9 at 4 dpi and HY03 at 11 dpi, so albeit that they represented a common pattern at the peak of innate immune response. COPD iALI responded earlier and stronger to SARS-CoV-2 infection with a common pattern to the 2 patients including overexpression of interferon-stimulated genes *MX1* at 1 dpi, *IFI35* and *IFITM3* 4 dpi, which are known cellular proteins involved in viral restriction (41–46).

Moreover, pro-inflammatory cytokine secretion induced by SARS-CoV-2 infection of iALI organoids was evaluated by measuring cytokines released in apical wash at 4 dpi with flow immunoassays (Supplemental Fig. 3). CCL5 and IL2 relative expression was found significantly higher in COPD iALI as compared to HY03 iALI. Interestingly, IL2 is one of the biomarkers in COPD patients to predict prognostic in respiratory failure (47). In addition, an increased concentration of G-CSF and CXCL10 was found released from infected HY03 and COPD iALI. Collectively, these results of iALI cytokine secretion in response to SARS-CoV-2 infection are significantly more pronounced in iCOPD as compared to healthy iALI.).

Overall, our results suggest that iALI bronchial organoids allowed for assessing individual innate immune response to infection. While our results are coherent with known SARS-CoV-2 induced ISG response, they revealed intrinsic differences between healthy and the two COPD patient response. Despite expected heterogeneities, further investigation could bring knowledge on COPD pathology and its response to viral infection for each individuals.

## Discussion

Our work shows that induced pluripotent stem cell derived bronchial organoids are an excellent bronchial epithelium model for reproducing several COPD characteristics, and for studying long lasting virus replication in bronchial tissue from healthy and COPD patients, opening doors for mechanistic studies and a better understanding of the contribution of genetic or epigenetic changes to the disease. As previously published (30), results confirm efficiency of our differentiation protocol of iPSC into functional airway epithelium. Nevertheless, we observed unexpected differences between organoids produced from HY03 and COPD iPSC. Indeed, reprogramming of somatic cells into iPSC is known to erase most of the epigenetics features to reset gene expression to a pluripotent state (48–50). Recent studies have nonetheless proven maintenance of some donor specific DNA methylation and gene expression in iPSC (51–54). In our work, iPSC were reprogrammed from blood cells and not airway cells that would most likely carry epigenetic marks induced by noxious particles exposition. So, we hypothesize that donors iCOPD2 and iCOPD9 have some genetic predisposition to COPD associated to impaired immune response to SARS-CoV-2 infection. Indeed, the two donors were considered light smokers compared to other COPD patients but still developed a severe form of COPD very early in life and reported a familial history of COPD (Supplemental Table 3).

SARS-CoV-2 successfully infected iALI and impacted the airway epithelium similarly to what has been previously reported, notably cell shedding (11), cilia disruption (21) and mucus secretion (55,56). The early innate immune response observed is also consistent with results described in primary models (10, 34, 35). We especially note expression of well-known antiviral restriction ISGs such as *OAS2, IFT1, IFITM3* and *MX1* (41–46). *ISG15, IFIT1, IFIT3, IFI44* and *CXCL10* were found over expressed in a primary air-liquid interface model (57). *MX1, ISG15, IFI44, IFITM1* were significantly increased in infected bronchioalveolar air-liquid interface model derived from fetal lung bud tip organoids (58). iPSC derived organoids reported overexpression of *OAS2, IFIT1, IFIT3, IFI44, MX1, ISG15* in the airway (59) and *IFITM1*and *IFI35* in the alveolus (36).

As mentioned, previous preclinical models of COPD were developed from lung progenitors and gave conflicting results as recently reported (10,34,35), notably a lower innate immune response in response to SARS-CoV-2 infection (10,35). However, one main limitation of these studies is the duration (3 to 4 dpi) over which they studied SARS-CoV-2 infection. In our iALI models, a thick layer of mesenchymal cells is present, supposedly allowing to maintain iALI viability and stability for more than three weeks. After 4 dpi and up to 18 dpi, our results show that SARS-CoV-2 infection persists and show differences in response to infection contrary to what has been previously reported. We observe some disparities between HY03 and COPD donors especially in the intensity of their innate immune response to SARS-CoV-2 infection. The variation of innate immune response correlates with the viral RNA measured in the iALI cells. At one day post-infection, it is striking that viral replication is significantly lower in COPD iALI than in healthy that do not show, yet, innate immune response. But at 4 days post-infection, iCOPD2 iALI shows highest viral replication and lowest innate immune response. In contrast, significantly lower intracellular viral replication was observed in iCOPD9 as compared to iCOPD2 iALI. Indeed, iCOPD9 shows the strongest innate immune response at 4 days post-infection. We thus observed an early and high innate interferon responses limiting viral replication in COPD iALI. In contrast, HY03 iALI donor only shows an innate immune response at 11 days post-infection, when the peak of viral infection has passed. Unfortunately, we were unable to analyze iCOPD9 iALI donors at 11 days post-infection to determine whether their innate immune response was sustaining further.

It is noteworthy that the level of mucus in iCOPD is already at a maximum level at 1dpi in contrario to healthy iALI that never reach this level even 11 dpi which can have some consequences on (1) sensing SARS-CoV-2 and (2) on the response to antiviral drugs.

Finally, a major limitation of viral infection response studies on iALI, similarly to studies on primary airway epithelium, is the absence of immune cells. To be more accurate, further development of the iALI model would include co-culture of immune cells that would necessitate to be differentiated from the same iPS, which promises to be limited and difficult to achieve.

Our work on iALI model revealed differences between our three donors, that modulated their response to airway infection. It is important, to note that these differences, hypothesis as genetic predisposition, could also be due to either heterogenous iPS reprogramming, or a difference in donor sex, âge and history. Conclusion on COPD disease would require to increase the number of donors to a significant amount that could not be achieved during that study.

Overall, our results indicate that these iALI react early to SARS-CoV-2 infection with a strong functional and interferon response, simiraly to literature data (11,21,36,41–46,55–58). The observed variability between donors could be linked to our previous hypothesis of individual genetic predisposition to COPD, that might impact differently the cells and therefore the inflammation and innate immune response of the tissue, despite the absence of immune cells. Nevertheless, our results show a higher and earlier innate immune response to SARS-CoV-2 in COPD samples, which correlates with clinical data reporting high susceptibility to a severe form of COVID-19 (4–7), and suggests that SARS-CoV-2 infection is more likely to trigger an early cytokine storm in patients with COPD, less response to antivirals and more chance to develop severe COVID-19.

## Materiel and Methods

### iALI

iPSC-derived airway epithelium on ALI (iALI model) was generated as previously described (27) from the hiPSC lines HY03 (UHOMi002-A) (healthy control), iCOPD2 (UHOMi003-A), and iCOPD9 (UHOMi005-A) (27,60). Briefly, the major stages of embryonic lung development were recapitulated as follows: stage 1, definitive endoderm (day 0–3) using activin A (100 ng/ml), Y-27632(10µM), CHIR-99021 (3Mm) between 16h and 40h post plating, and LDN-193189 (250nM) between 40h and 70h post plating; stage 2, anterior foregut endoderm (day 4–7); stage 3, after transfer into a transwell, lung progenitor specification (day 7-12); stage 4, polarized epithelial layer (day 12-40), and day 40+: multi-ciliated bronchial epithelial layer. After 40 days, the airway epithelium on iALI displays morphologic and functional similarities with primary human airway epithelial cells and included different airway cell types (basal, secretory, and multi-ciliated cells).

### SARS-CoV-2 virus stock and titration

The strains Delta (B.1.617.2) hCoV-19/France/HDF-IPP11602i/2021 were supplied by the National Reference Centre for Respiratory Viruses hosted by Institut Pasteur (Paris, France) and headed by Pr. Sylvie van der Werf. The human sample from which strain hCoV-19/France/HDF-IPP11602i/2021 was isolated has been provided by Dr Guiheneuf Raphaël, CH Simone Veil, Beauvais France. Moreover, the strain was supplied through the European Virus Archive goes Global (EVAg) platform, a project that has received funding from the European Union’s Horizon 2020 research and innovation program under the grant agreement No 653316. The viruses were propagated in VeroE6 cells with DMEM containing 25mM HEPES at 37°C with 5% CO2 and were harvested 72 hours post inoculation. Virus stocks were stored at −80°C. Virus titration was monitored using plaque assay.

### Viral infection

iALI cultures were washed 24 h before infection by adding culture medium to the apical side at 37 ◦C for 15 min. For SARS-CoV-2 infection, iALI apical side was covered with 350 µL of viral dilution in serum free DMEM (2,5 × 10^5^ PFU per sample, estimated MOI 0,05) for 1 h 30 min at 37 ◦C. For remdesivir efficiency evaluation infection was done on smaller iALI at an estimated MOI 0,01 (0,5 × 10^5^ PFU per sample) Then, the inoculum was removed, and the culture was quickly washed with culture medium PneumaCult-ALI Medium (05001, Stem cell technologies). The first point of the kinetic analysis (Day 0) was collected by adding culture medium to the apical side at 37◦C for 15 min. Some iALI’s apical side were washed everyday with culture medium to collect kinetics samples. Other iALI were washed only at the time of collection (RLT or 4% PFA). SARS-CoV-2 infection experiments were conducted inside the BSL3 facility at the CEMIPAI institute.

### Reverse Transcription-Quantitative Polymerase Chain Reaction (RT-qPCR)

**Cellular RNA** of iALI were collected in RLT and extracted using the QIAshredder kit (QIAGEN, Redwood city, CA, USA) and the RNeasy mini kit (Qiagen, Redwood city, CA, USA) according to the manufacturer’s instructions.

Gene expression was quantified by RT-qPCR in triplicate using the Luna Universal One-Step RT-qPCR Kit (New England Biolabs, Ipswich, MA, USA) and a BIORAD CFX Opus 384 system. Primers are described in Supplemental table 1. Relative gene expression was calculated by normalizing to GAPDH gene level or GAPDH and Epcam gene level (control) and using the ΔCt method. Cell type markers expression are displayed as relative expression on log2 scale. The gene expression of the innate immune response is presented as the difference from the log2 fold change gene median, with values depicted according to the color scale shown on the right.

Viral RNA was extracted of daily apical wash with NucleoSpin Dx Virus (Macherey-Nagel). Absolute quantification of the viral envelope E gene was done, in triplicate, via RT-qPCR as described above and by comparison to standard range. Data are presented as the concentration of viral RNA in the apical wash.

### Samples embedding and cross section

After collecting the apical wash, the iALI were quickly rinsed with PBS and fixed in 4% paraformaldehyde for 24 hours at 4°C. The samples were then embedded in 5% agarose and placed in 70% ethanol before being sent to the Montpellier histology platform (RHEM). RHEM services included paraffin embedding, 3-5µm cross-sections, hematoxylin-eosin-safran (HES) staining, and periodic acid-Schiff-blue alcian (PAS-BA) staining. The stained slides were scanned using a NanoZoomer, Hamamatsu, by the Montpellier Imagery Platform (MRI).

### Cross-section PAS-BA staining analysis

Using PAS-BAstained slices, the sample and mucus cell areas were identified and analyzed with Histometrix software (Quantacell). Detection was carried out using two U-Net deep learning models trained on a selected subset of the dataset. For ensuring reliability, the results were subsequently verified through visual inspection. All analysis was done by Quantacell, Montpellier, France.

### Immunofluorescence

Immunofluorescence was used to label the cross-section’s slides (CS) and unprocessed/cut organoids. Fixation of organoids for direct immunofluorescence was done by immersion in 4% paraformaldehyde for 30 minutes at room temperature.

For cross-section slides, preliminary steps were needed to remove paraffin: A series of three-minute baths in xylene (*28973.294*, VWR), ethanol 100%, ethanol 96%, ethanol 70%, and ethanol 50% were used for rehydrating the samples. Slides were then washed with cold water and submerged for 20 minutes in sodium citrate pH 6 buffer (C9999, Sigma) at 100°C.

Following a permeabilization period of 20-minute for CS and 2h for organoids with PBS 0.5% triton, the slides were blocked for one hour for CS and two hours for organoid using blocking buffer (1% BSA, 0.1% Triton X-100, 10% donkey serum in PBS). Samples were incubated with primary antibodies (Supplemental table 2) with appropriate dilutions in staining buffer (1% BSA, 0.1% Triton X-100, in PBS) for overnight at 4°C, followed by washing three times with washing buffer (1% BSA, 0.025% Triton X-100 in PBS). AlexaFluor 488, 555, or 647-labeled secondary antibodies two hours incubation at room temperature followed and then three times wash with washing buffer. Then, 4′,6-diamidino-2-phenylindole (DAPI) was incubated for a brief 5 minutes for CS and 15 minutes for organoids and washed three times with PBS washing buffer. Finally, aqueous mounting media (BUF058B, Biorad) was used to cover CS with a glass coverslip and mount organoids on glass slides, and glass coverslips were used to cover them. The Cell Discovery 7 LSM900 Airyscan2 microscope was used to perform high resolution 3D imaging, and ZEN Image analysis software (Zeiss) was used for processing. All of the primary antibodies utilized in this investigation were confirmed, and information about them is included in Supplementary Table xx. All quantitatively analysis were done using ZEN Image analysis software (Zeiss) and Fiji/Image J software.

### Plaque assay

Virus titration from stocks and infected cell culture supernatants were monitored using plaque assays on a monolayer of VeroE6 cells, using 100µL of serially diluted samples. The plaque forming unit (PFU) values were determined using crystal violet coloration on cells and subsequent scoring the wells displaying cytopathic effects. Calculations allow to determine the titer as the number of PFU/mL.

### Immunoassays

Cytokine quantification was done on 4dpi everyday apical wash with the LEGENDplex COVID-19 Cytokine Storm Panel 1 (14-plex) with V-bottom Plate (741089, Biolegend) samples were measured at 2 dilutions, pure and 10 fold dilution to cover the standard range of all the analytes. Acquisition was done on the NovoCyte Flow 21000YB cytometer inside the CEMIPAI BSL3 facility. Results were normalized to total protein levels as measured by pierce protein assay (10177723, thermo fisher).

### Statistical analysis

For normally distributed data, the mean is shown with standard error; where necessary, an analysis of variance (ANOVA) or Student t test is employed to look for intergroup differences. *P <0.05, **P <0.01, ***P <0.001, and ****P <0.0001 demonstrate the degree of significance, with a p-value <0.05 being considered significant. Graphical representation and statistical analysis were performed using GraphPad Prism

## Acknowledgments

We are grateful to CEMIPAI UAR3725 CNRS/University of Montpellier, France for supporting BSL3 facility and Microscopy and for excellent technical services. We thank Carine Bourdais, Agathe Cœur and Cecilia Urena from IRMB for iALI culture technical advices and their much-needed help for maintenance over week-ends. We also thanks Romane Pisteur, master student, who draw the epithelium scheme.

We acknowledge the “Réseau d’Histologie Expérimentale de Montpellier” -RHEM facility for histology technics and expertise. RHEM facility is supported by SIRIC Montpellier Cancer Grant INCa_Inserm_DGOS_12553, REACT-EU (Recovery Assistance for Cohesion and the Territories of Europe), IBiSA, Ligue contre le cancer, the Occitanie / Pyrénées-Méditerranée and GIS FC3R whose funds are managed by Inserm.

## Authors contributions

D.M and J.D.V. conceptualized the study; D.M, J.D.V., NG and SA supervised the study. N.G. did viral production and titration. L.M., F.F., A.N. cultured iPSc cells and organoids. L.M. performed viral infection on iALI organoids and RTqPCR, virus collect and titration, immunoassays and immunofluorescence on iALI, and analyse data. J.S. performed confocal imaging and quantitative image analysis. G.G. and V.R. performed FFPE staining image analysis. L.M and DM wrote the manuscript. N.G., S. A., A.B. and J.D.V. contributed to manuscript editing.

