## Supplemental material for "Human pluripotent stem cell-derived bronchial airway organoids provide insights into differential innate immune and long-term responses to SARS-CoV-2 infection in healthy and COPD"

^1^ CEMIPAI: Centre d’Etudes des Maladies Infectieuses et Pharmacologie Anti-Infectieuses, CNRS UAR3725, ^2^ IRIM UMR 9004, CNRS & Université de Montpellier - ^3^ IRMB : Institute of Regenerative Medecine and Biotherapy, Université de Montpellier, CHU de Montpellier, INSERMU1183, ^4^ QuantaCell, IRMB, CHU de Montpellier, 80 Av. Augustin Fliche, 34090 Montpellier. ^5^ Department of Respiratory Diseases, CHU Montpellier, Arnaud de Villeneuve Hospital, INSERM, Montpellier 34000, France; PhyMedExp, University of Montpellier, INSERM U1046, CNRS UMR 9214.

**Correspondence:** **Co-last senior authors,* *(DM);* *(JDV)*

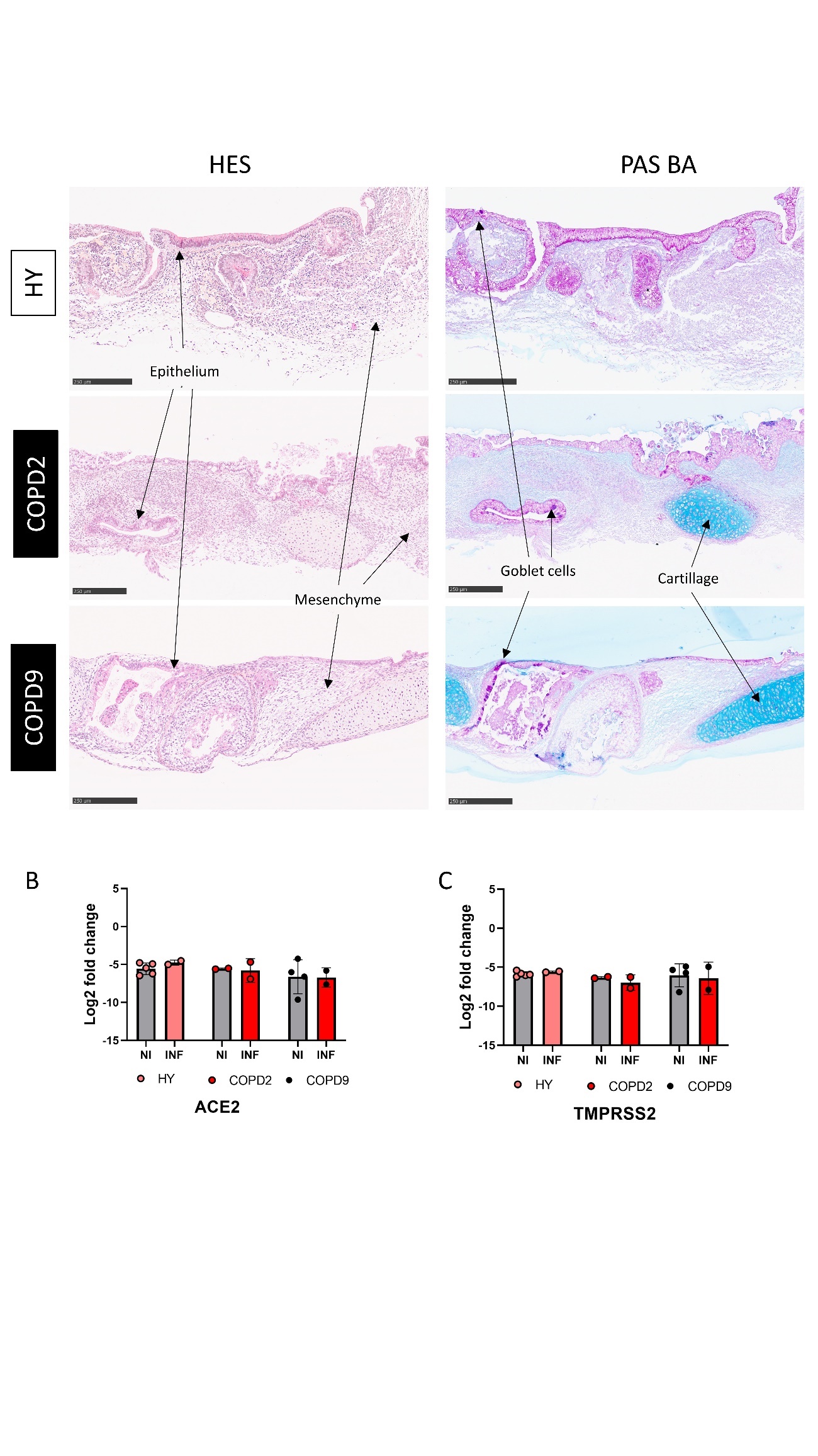

(A) **Supplemental Figure 1: Imaging of stained iALI cross sections.** Representative hematoxyline eosine safran (HES) and periodic acid Schiff – blue alcian (PAS-BA) stained cross sections of HY03 and iCOPD iALI; (B) ACE2 and (C) TMPRSS2 relative expression to GAPDH in non-infected and infected HY03, iCOPD2 and iCOPD9 iALI measured by RTqPCR. NI: non infected; INF: SARS-CoV-2 infected.

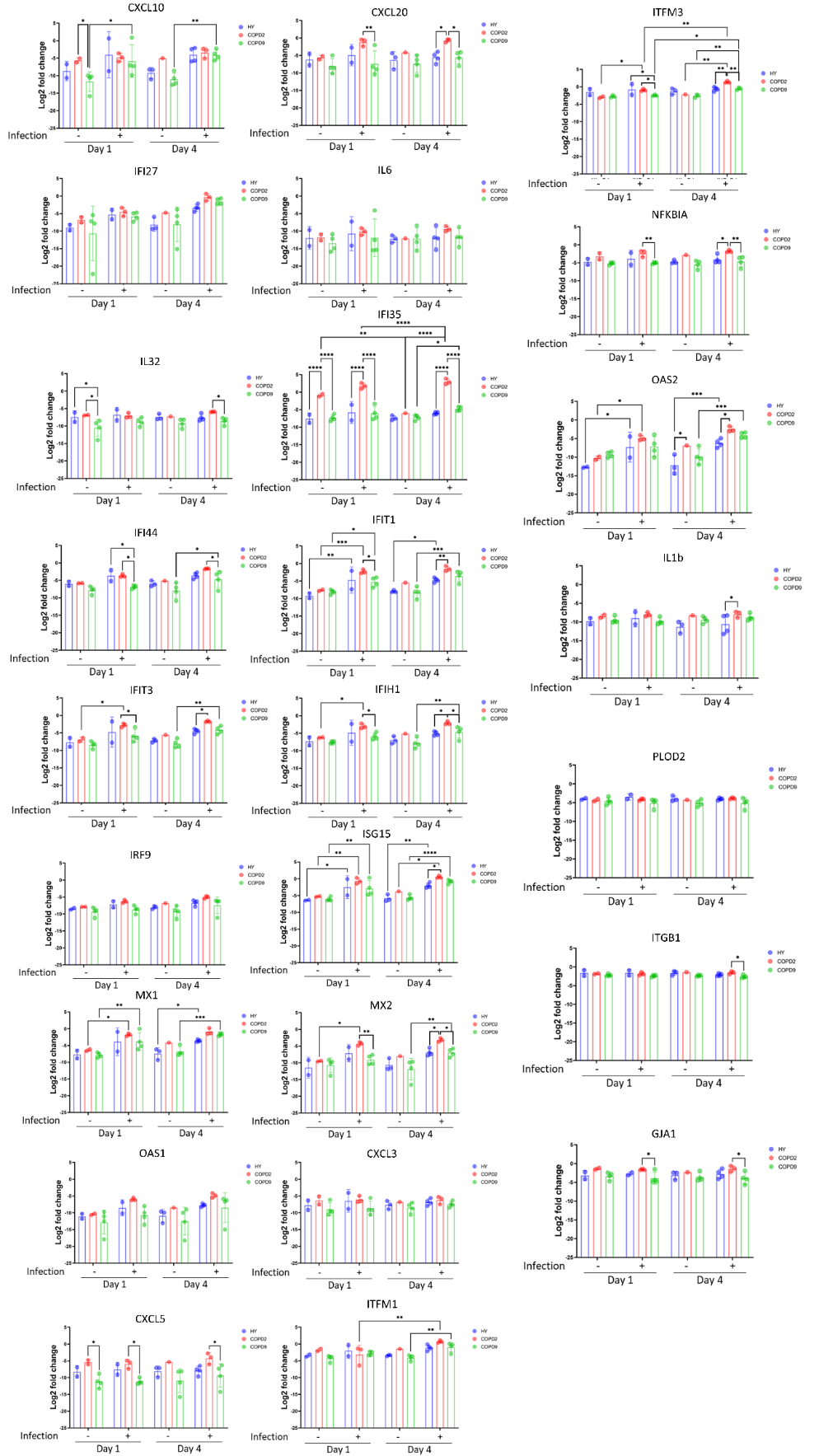

**Supplemental Figure 2: Innate immune response in HY03 and iCOPD iALI 1 and 4 dpi .**

Mean innate immune response gene expression measured by RTqPCR in non infected HY03 (n=2), iCOPD2 (n=2) and iCOPD9 (n=4) iALI, 1 dpi in HY03(n=2), iCOPD2 (n=3) and iCOPD9 (n=4) iALI, or, 4 dpi in HY03 (n=4), iCOPD2 (n=3) and iCOPD9 (n=4) iALI with the non infected counterparts. Data are presented as the mean of the log 2 relative expression to GAPDH for each condition. *p < 0.05, **p < 0.01, ***p < 0.001, **** p < 0.0001, NS represents not significant, two-way ANOVA with Tukey’s multiple comparisons post-test N= Number of experiments, n= total number of samples analyzed.

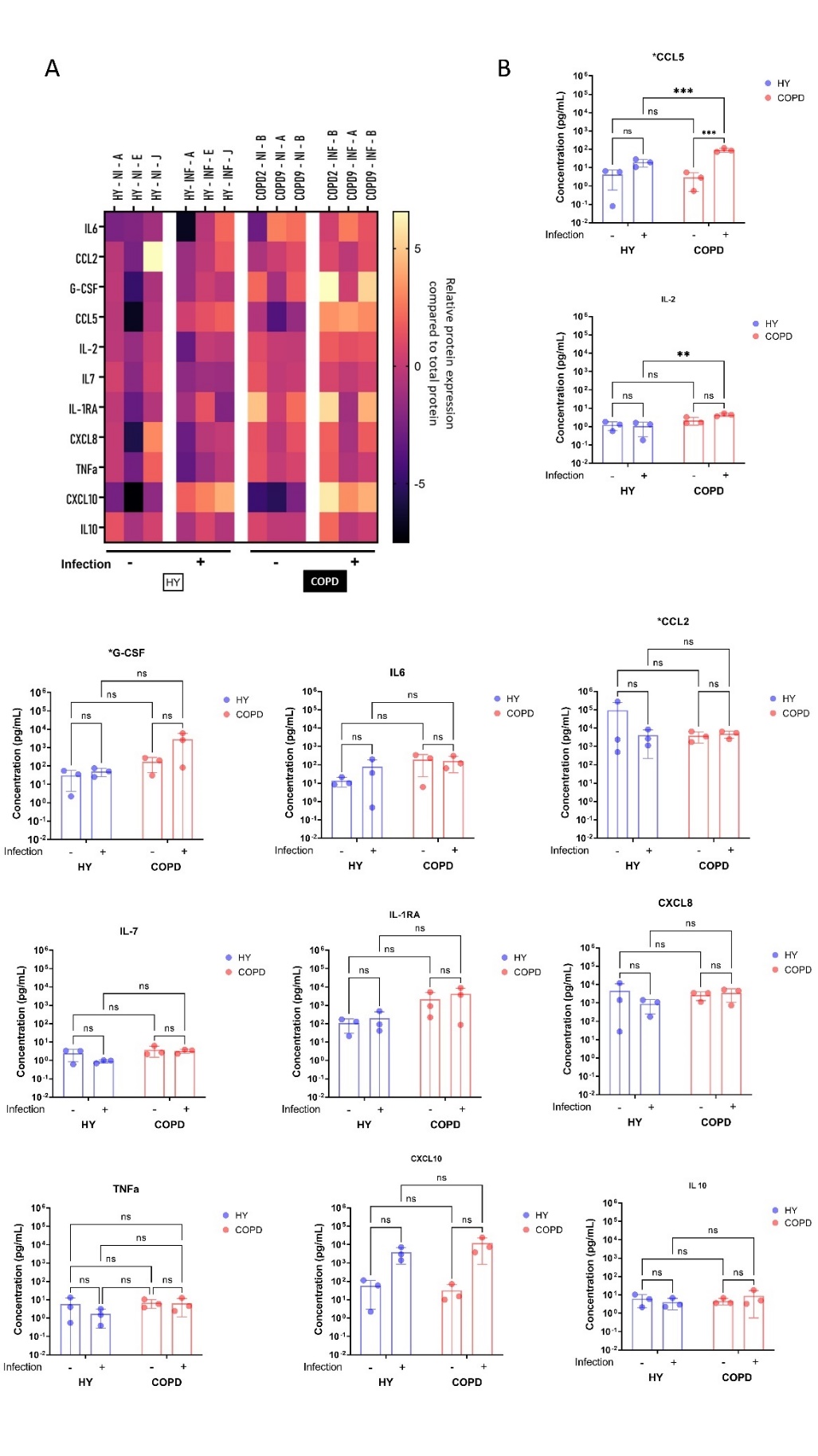

**Supplemental Figure S3: Proteomic analysis of innate immune response of SARS-CoV-2 infected iALI organoids.** Inflammation and innate immune response in HY03 and COPD iALI was measured at protein level by multi-analyte flow assay targeting COVID-19 cytokine storm effector (IL-6, CCL2, G-CSF, CCL5, IL-2, IL7, CXCL8, TNFα, CXCL10, IL10) in non-infected or infected (4dpi). (A) Data are presented with a heat map where each value represents the difference from the median of the log2 protein concentration normalized to total protein concentration and is depicted according to the color scale shown at the right (-5 to 5). (B) Mean innate immune response protein concentration measured by legend-plex and normalized to total protein concentration measured by BCA assay 4 dpi or non infected in HY03(n=3), iCOPD2 (n=1) and iCOPD9 (n=2) iALI. Data are presented as the mean of the normalized protein concentration. *p < 0.05, **p < 0.01, ***p < 0.001, **** p < 0.0001, NS represents not significant, two-way ANOVA with Sidak’s multiple comparisons post-test N= Number of experiments, n= total number of samples analyzed. Each experiment is coded with an alphabet letter (ie., A to J). NI: non infected; INF: SARS-CoV-2 infected.

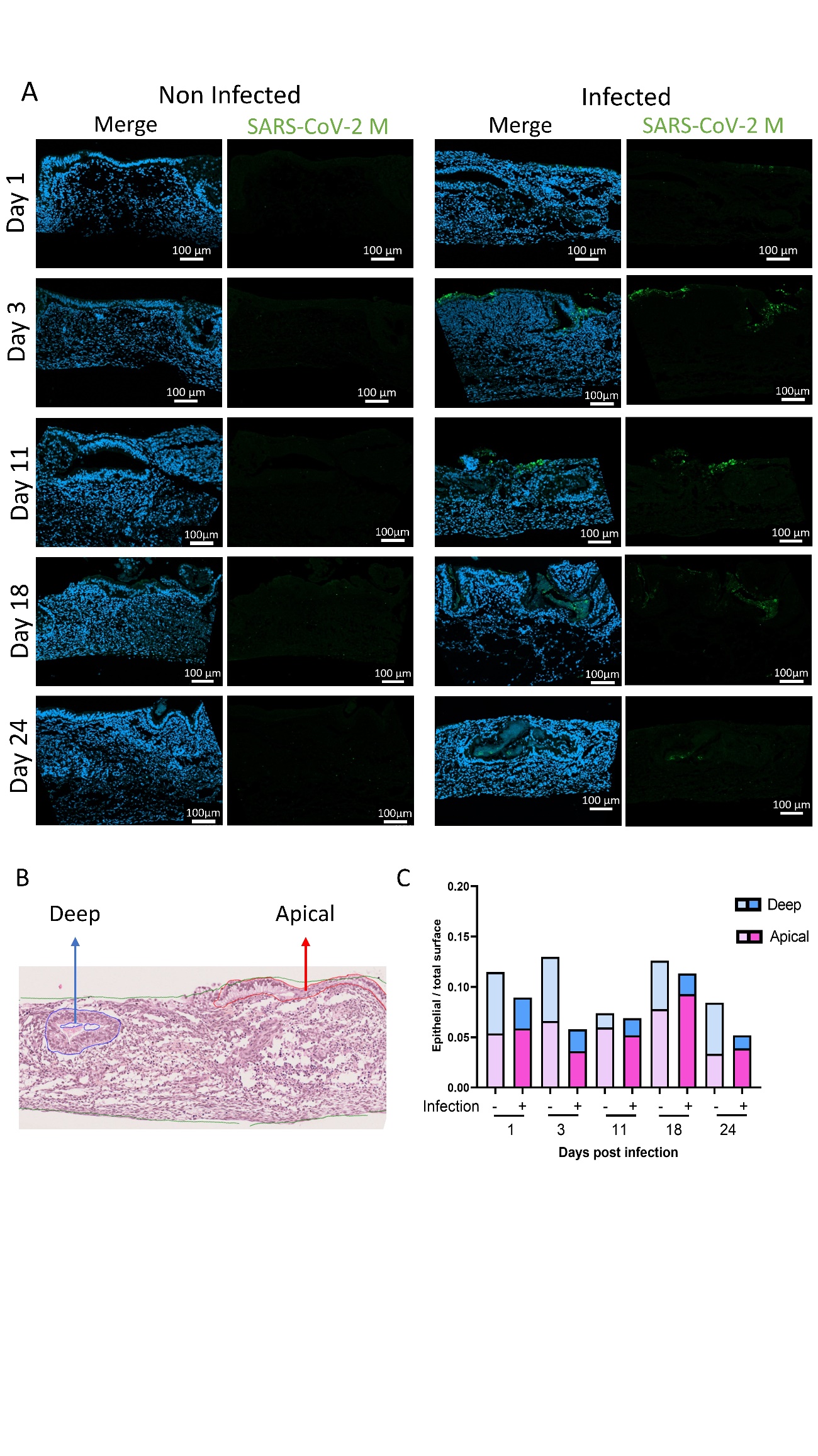

**Supplemental Figure S4: SARS-CoV-2 infection in iALI tissue overtime.**

(A)SARS-CoV-2 infection was visualized by labelling SARS-CoV-2 membrane protein (Virus M, green) in healthy iALI (cross section). (B) Example of HES labelling and segmentation of apical and deep bronchial epithelium. (C) Quantification of the area of apical and deep epithelial tissue normalized to total sample area on non infected and infected iALI 1, 3, 11, 18 and 24 dpi.

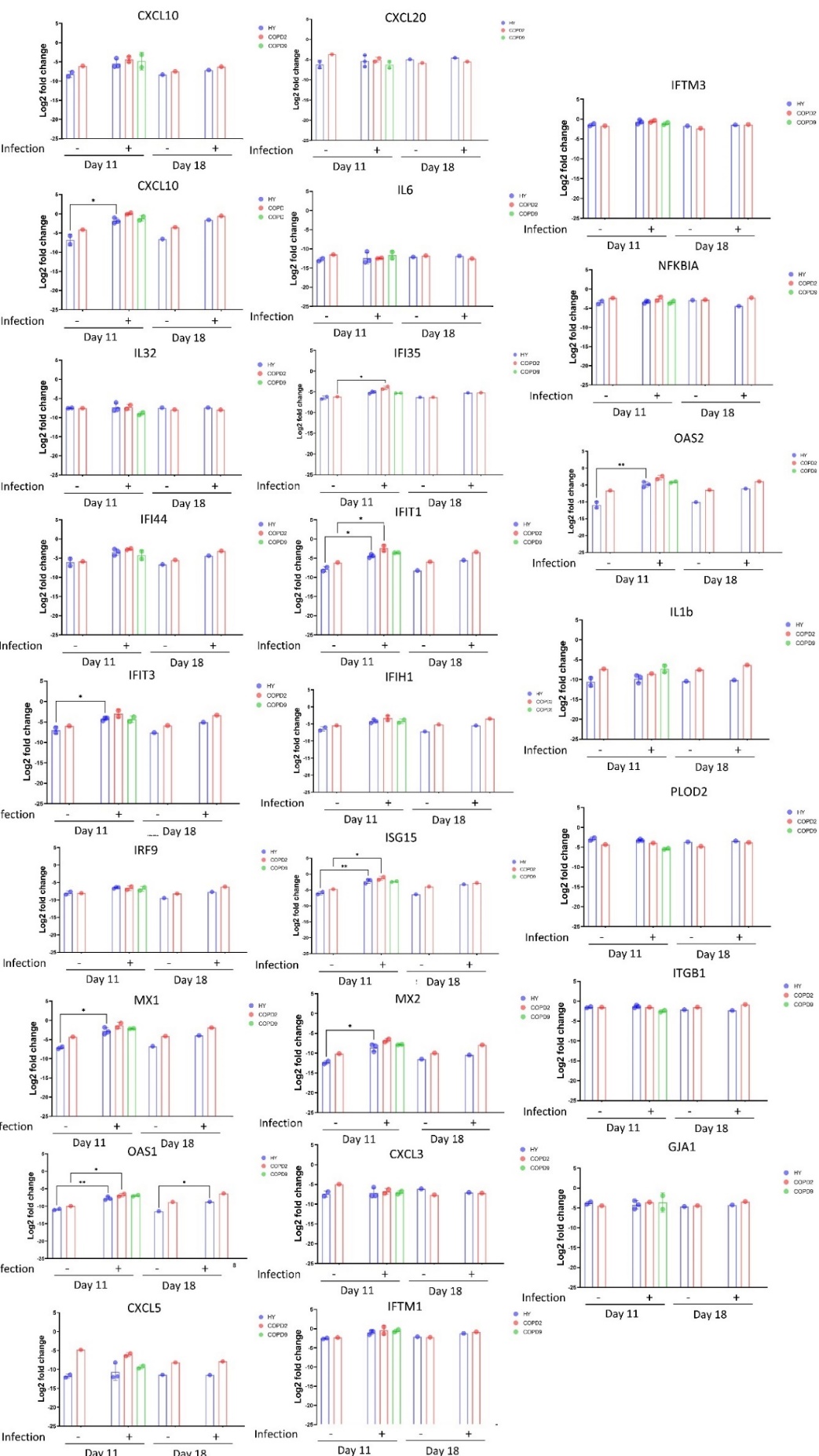

**Supplemental Figure S5: Innate immune response in HY03 and COPD iALI 11 and 18 dpi.**

Mean innate immune response gene expression measured by RTqPCR in non infected HY03(n=2) and iCOPD2 (n=2) iALI, or 11 dpi in HY03(n=3), iCOPD2 (n=2) and iCOPD9 (n=2) iALI, 18 days non infected HY03 (n=1) and iCOPD2 (n=1) iALI, or 18 dpi in HY03 (n=1) and iCOPD2 (n=1) iALI. Data are presented as the mean of the log 2 relative expression to GAPDH for each condition. *p < 0.05, **p < 0.01, ***p < 0.001, **** p < 0.0001, NS represents not significant, two-way ANOVA with Tukey’s multiple comparisons post-test N= Number of experiments, n= total number of samples analyzed.

|  | **Primer forward** | **Primer reverse** |
| --- | --- | --- |
| FOXJ1 | ACTCGTATGCCACGCTCATCTG | GAGACAGGTTGTGGCGGATTGA |
| CCDC40 | CCAGCAGTGGGCAGATTGA | CCAGGTCTGAGGACTCGATGT |
| Muc5AC | CCACTGGTTCTATGGCAACACC | GCCGAAGTCCAGGCTGTGCG |
| Muc5B | CTGCTACGACAAGGACGGAAA | AAGGCTGTGAGCGCACTGGATG |
| KRT5 | GCTGCCTACATGAACAAGGTGG | ATGGAGAGGACCACTGAGGTGT |
| P63 | CGCCATGCCTGTTCACAA | ACTGCCAAATTGCAAAGACA |
| CHGA | CGGATCCTTTCCATTCTGAG | ACCGCTGTGTTTCTTCTGCT |
| SCGB1A1 | GCTGAAGAAGCTGGTGGACACC | GCGTGGACTCAAAGCATGGCAG |
| GAPDH | GAC CTG ACC TGC CGT CTA GAA A | CCT GCT TCA CCA CCT TCT TGA |
| CXCL10 | CCTCCAGTCTCAGCACCATG | TGAAGCAGGGTCAGAACATCC |
| CXCL20 | CCTCTGCGGCGAATC AGAAG | CTGCCGTGTGAAGCCCACAA |
| IL6 | AGACAGCCACTCACCTCTTCAG | TTCTGCCAGTGCCTCTTTGCTG |
| IL32 | GACCTCTGTCTCTCTCGGGT | CTGTCTCCAGGTAGCCCTCT |
| IFI27 | CTTCACTGCGGCGGGAATC | CCAGGATgAACTTGGTCAATCC |
| IFI35 | GAGGGTGTTGGTCACTGGATT | CA CCTCCGTTCCTAGTCTTGC |
| IFI44 | TCCAAGGGCATGTAACGCAT | CCTCCCTTAGATTCCCTATTTGCT |
| IFIT1 | GCCTTGCTGAAGTGTGGAGGAA | ATCCAGGCGATAGGCAGAGATC |
| IFIT3 | GAAACAGCCATCATGAGTGAGG | GCATCTGAGAGTCTGCCCAA |
| IFIH1 | GAACCTCCTTCAGCCCACTC | CCCATGGTGCCTGAATCACT |
| ISG15 | GCAGATCACCCAGAAGATCG | GGCCCTTGTTATTCCTCACC |
| IRF9 | TATCAGTTGCTGCCACCAGG | TAGGATGCCCCTCTCAAGCT |
| OAS1 | CCTACCCTGTGTGTGTGTCC | GCATAGACCGTCAGGAGCTC |
| MX1 | GGCTGTTTACCAGACTCCGACA | CACAAAGCCTGGCAGCTCTCTA |
| MX2 | GCGATGCCTGTGGTTGTTTT | GAGTGGAGAGGCTGGGAAAC |
| ITFM1 | TCGCCTACTCCGTGAAGTCTA | TGTCACAGAGCCGAATACCAG |
| ITFM3 | GGTCTTCGCTGGACACCAT | TGTCCCTAGACTTCACGGAGTA |
| CXCL3 | AACCGAAGTCATAGCCACAC | TGCTCCCCTTGTTCAGTATC |
| NFKBIA | CTC CGA GAC TTT CGA GGA AAT AC | GCC ATT GAA GTT GGT AGC CTT CA |
| OAS2 | CTGGAGCTGGTCACACAATATC | GAAACTTCCTCACGGTCTCATC |
| CXCL5 | TGTGTTGAGAGAGCTGCGTT | CCAGTGATTCCTGGCTCACA |
| PLOD2 | ACTCCCCTACTCCGGAAACA | TGAGGACGAAGAGAACGCTG |
| IL1b | CCACAGACCTTCCAGGAGAATG | GTGCAGTTCAGTGATCGTACAGG |
| ITGB1 | TGATTGGCTGGAGGAATGTTA | GTTTCTGGACAAGGTGAGCAA |
| GJA1 | GGCCTTCTTGCTGATCCAGT | GCTGGTCCACAATGGCTAGT |
| Enveloppe viral | ACAGGTACGTTAATAGTTAATAGCGT | ATA TTG CAG CAG TAC GCA CAC A |
| Epcam | GCCAGTGTACTTCAGTTGGTGC | CCCTTCAGGTTTTGCTCTTCTCC |

**Supplemental Table S1: Primer sequence**

| **Primary antibody** | **Reference** | **Dilution** |
| --- | --- | --- |
| CDHR3 | HPA011218, Sigma | 1/100 |
| Muc5Ac | ab3649, Abcam | 1/100 |
| KRT5 | PA1-37974, Thermofisher | 1/100 |
| P63 | AF1916, R&D systems | 1/200 |
| CC10 | RD181022220-01, Biovendor | 1/1000 |
| TubIV | T7941, Sigma | 1/200 |
| SARS-CoV-2 M | PAB31758, Abnova | 1/200 |
| DAPI | D9542, Sigma | 1/2000 |
| **Secondary antibody** | **Reference** | **Dilution** |
| Donkey anti-Rabbit IgG – Alexa Fluor 488 | A21206, Invitrogen | 1/1000 |
| Donkey anti-Mouse IgG – Alexa Fluor 555 | A31570, Invitrogen | 1/1000 |
| Donkey anti-Goat IgG – Alexa Fluor 647 | A21447, Invitrogen | 1/1000 |
| Donkey anti-Rabbit IgG – Alexa Fluor 647 | ab150075, abcam | 1/500 |
| Donkey anti-Goat IgG – Alexa Fluor 488 | ab150129, abcam | 1/1000 |

**Supplemental Table S2: Antibodies references and dilutions**

|  | **HY03** | **iCOPD2** | **iCOPD9** |
| --- | --- | --- | --- |
| **Age** | 42 | 52 | 54 |
| **Sex** | M | F | M |
| **Body-mass index (kg/m²)** | 24 | 30 | 32 |
| **Smoking current/Ex-smoker** | No | Current | Ex-smoker |
| **Pack-year history** | N.A. | 45 | 75 |
| **Age of beginning (years)** | N.A. | 14 | 24 |
| **Others toxic** | No | Yes | No |
| **Pulmonary function and symptoms** | N.A. |  |  |
| **Age of diagnosis** |  | 48 | 49 |
| **Age at onset of first symptoms (years)** |  | 40 | 35 |
| **In utero smoking exposure** |  | Yes | Yes |
| **Premature birth** |  | No | Yes |
| **Familial history of COPD/emphysema** |  | Yes | Yes |

**Supplemental Table S3: PBMC donors characteristics**
